# Multilevel human secondary lymphoid immune system compartmentalization revealed by complementary multiplexing and mass spectrometry imaging approaches

**DOI:** 10.1101/2022.11.01.514691

**Authors:** Benjamin L. Oyler, Jeferson A. Valencia-Dávila, Eirini Moysi, Adam Molyvdas, Kalliopi Ioannidou, Kylie March, David Ambrozak, Laurence de Leval, Giulia Fabozzi, Amina S. Woods, Richard A. Koup, Constantinos Petrovas

**Affiliations:** Tissue Analysis Core, Immunology Laboratory, Vaccine Research Center, NIAID, NIH, Bethesda, MD, USA; Department of Laboratory Medicine and Pathology, Institute of Pathology, Lausanne University Hospital and Lausanne University, Lausanne, Switzerland

**Keywords:** tonsils, follicles, confocal imaging, mass spectrometry imaging

## Abstract

Secondary human lymphoid tissue immune reactions take place in a highly coordinated environment with compartmentalization representing a fundamental feature of this organization. *In situ* profiling methodologies are indispensable for the understanding of this compartmentalization. Here, we propose a complementary experimental approach aiming to reveal different aspects of this process. The analysis of human tonsils, using a combination of single cell phenotypic analysis based on flow cytometry and multiplex imaging and mass spectrometry-based methodologies, revealed a compartmentalized organization at cellular and molecular level. More specifically, the skewed distribution of highly specialized immune cell subsets and relevant soluble mediators was accompanied by a compartmentalized localization of several lipids across different anatomical areas of the tonsillar tissue. The performance of such combinatorial experimental approaches could lead to the identification of novel *in situ* interactions and molecular targets for the *in vivo* manipulation of lymphoid organ, particularly the germinal center, immune reactions.

## Introduction

Secondary lymphoid organs (SLO) like tonsils, lymph nodes and spleen are anatomical sites where the development of adaptive immunity to pathogens and immunogens takes place. The organization of these organs is characterized by the presence of areas well-defined by their specific stromal and immune cell phenotypes and distinct roles in the development of humoral responses (1). These areas have been extensively investigated previously at a cellular and molecular level (2–4), using animal models and several analytical methodologies. However, the spatial organization of relevant *in situ* dynamics are less understood in humans. Follicles and Germinal centers (GCs), paracortical area, T cell zones and medulla are major anatomical areas of lymphoid organs (5). Trafficking of pathogens or immunogens into lymphoid organs results in the initiation of a cascade of immune reactions, ultimately leading to the development of antigen-specific B cell responses. In the T-cell zone, the coordinated interaction between stromal cells (*e.g*. Fibroblastic Reticular Cells, FRC), innate immunity (monocytes, macrophages, dendritic cells), and naïve, non-differentiated CD4 T cells will start a unique reprogramming of CD4 T cells and interaction with B cells, ultimately leading to their differentiation and trafficking into the follicular areas (6–9). Within the follicle/GC, the orchestrated spatial organization and function of highly differentiated immune cell populations and stromal cells (*e.g*. Follicular Dendritic Cells, FDC) will lead to the development of antigen-specific B cell responses (10). Specifically, the GC is characterized by two distinct areas (11); the Light Zone (LZ), where the interaction between follicular helper T cells (T_FH_), GC B cells, antigen and FDC takes place (12, 13), and the Dark Zone (DZ), wherein the division and further maturation of B cells occurs (14). The outcome of the B cell responses is critically dependent on the quality and specificity of the “help” received from T_FH_ (15).

Compartmentalization of the relevant immune reactions and players is a critical component for their coordinated function. This compartmentalization could facilitate the ordered, multiphase development of these immune interactions while it likely provides checkpoints for their regulation and the suppression of unwanted immune reactivity like autoimmunity. The relative accessibility, compared to other human lymphoid organs, the abundance of relevant immune cell types and the well-preserved anatomical areas (*e.g*. GCs) makes reactive human tonsils a prototype lymphoid organ for the study of follicular immune dynamics with respect to local microenvironment. Here, we have applied complementary single cell analysis and imaging methodologies to further characterize the organization of tonsillar anatomy both at cellular and molecular levels. Our data provide an experimental approach for the study of the spatial organization within human lymphoid organs and the identification of cellular and molecular signatures associated with micro-anatomical tissue structures. Furthermore, we propose a novel approach for the investigation of tissue organization using lipid signatures while a discussion of lymphocyte lipid composition is also provided.

## Materials and methods

### Human material

Tonsils were anonymized discarded pathologic specimens obtained from Children’s National Medical Center (CNMC) under the auspices of the Basic Science Core of the District of Columbia Developmental Center for AIDS Research. The CNMC Institutional Review Board determined that study of anonymized discarded tissues did not constitute ‘human subjects research’. Upon receipt, tissues were washed with ice-cold medium R-10 (RPMI 1640 supplemented with 10% fetal bovine serum, 2 mM L-glutamine, 100 U/mL penicillin and 100 μg/mL streptomycin, Invitrogen). Following the removal of surrounding fatty tissue and necrotic areas, part of the tissue was dedicated for fixation (formalin) and paraffin embedded block or snap frozen block preparation while the remaining tissue was cut into small pieces for cell preparation by mechanical disruption, followed by Ficoll-Paque density gradient centrifugation. Cells were stored in liquid nitrogen until further use.

### Flow cytometry studies

The following titrated, conjugated antibodies were used for flow cytometry analysis: CD3-H7APC (SK7, BD Biosciences), CD4-BV650 (SK3, BD Biosciences), CD8-Pacific Blue (RPA-T8, BD Pharmingen), CD19-FITC (J3-119, Beckman-Coulter), CD27-BV605 (O323, Biolegend), CD45RO-Cy5PE (UCHL1, BD Pharmingen), PD-1-BV711 (EH12.2H7, Biolegend), CD57-AF594 (NK-1, Novus Bio), IgD-PE (Goat polyclonal, Southern Biotech), CD20-BV570 (2H7, Biolegend), CD38-BV786 (HIT2, Biolegend). 1-2×10^6^ tonsil-derived cells were thawed and rested for 2h in a cell culture incubator before staining. Cells were washed with PBS, BSA (0.5%), incubated (5 min, RT) with a viability dye (Aqua-dye, Invitrogen) and surface stained with titrated amounts of antibodies for 30 min at room temperature. After washing, cells were resuspended and fixed with 1% paraformaldehyde. ***Data acquisition and analysis:*** 0.5-1×10^6^ events were collected on a BD LSR Fortessa X-50 flow cytometer (BD Immunocytometry Systems). Electronic compensation was performed with antibody capture beads (BD Biosciences). Data were analyzed using FlowJo Version 9.9.4 (Tree Star). Forward scatter area vs. forward scatter height was used to gate out cell aggregates.

### Immunofluorescence microscopy studies

#### Confocal microscopy imaging

FFPE tissue sections of 8 μm were prepared using a microtome (Leica) and processed (heat drying, deparaffinization) for antigen retrieval (Borg RTU, Biocare Medical) and antibody staining using titrated antibodies as recently described (16, 17). The following combination of antibodies was used: IgD (primary rabbit, clone EPR6146, abcam), CD8 (primary mouse IgG2b, clone 4B11, Invitrogen), CD68 (primary mouse IgG1, clone KP-1, DAKO), CD20-eF615 (clone L26, eBioscience), Ki67-AF647 (clone B56, BD Horizon), CD4-AF700 (goat polyclonal, R&D), PD-1-AF488 (goat polyclonal, R&D), CD57-BV480 (clone NK1, BD Horizon), and JoJo (nuclear marker, Life Technologies). The following titrated secondary antibodies were used: donkey anti-rabbit Brilliant Violet 421 (poly 4064 Biolegend), goat anti-mouse IgG2b Alexa Fluor 546 (A21143, Thermo Scientific) and goat anti-mouse IgG1 Alexa Fluor 546 (A-21123, Thermo Scientific). ***Data acquisition and analysis***: stained slides were imaged on a NIKON (A1) inverted confocal microscope system, equipped with 40X, 1.3 NA oil objective lens. Tissue sections stained with single fluorophores were used for the correction of the spectral spillover between channels through live spectral unmixing in NIS software. Image acquisition was performed with NIS-elements software and the data analyzed with Imaris software version 9.6 (Bitplane). Histocytometry analysis was performed as previously described (17). Briefly, imaging datasets were segmented post-acquisition based on nuclear staining and average voxel intensities for all channels were extrapolated in Imaris. Channel statistics were exported to csv (comma separated values) files format and analyzed in FlowJo version 10.6.1.

#### Multispectral imaging

FFPE tissue sections of 4 μm were cut and prepared for immunofluorescence staining using the following combination of antibodies against CD14 (clone EPR3653, Cell Marque, detection by Opal 570), FDC (clone CNA.42, Invitrogen, detection with Opal 690), CXCL-13 (rabbit polyclonal, ThermoFisher, detection with Opal 520), IL-17 (goat polyclonal, R&D systems, detection with Opal 620), CD20 (clone L26, ThermoFisher, detection with Opal 480) and DAPI (nuclear marker), as previously described (18). Briefly, the staining consisted of consecutive rounds of antigen retrieval, staining with primary antibody, secondary HRP-labeled antibody and detection with optimized fluorescent Opal tyramide signal amplification (TSA) dye (Opal 7-color Automation IHC kit, from Akoya, Ref. NEL821001KT) and repeated antibody denaturation cycles. Multispectral images were acquired using the latest Vectra Polaris imaging system from Akoya. All images were recorded using a 20x magnification. The Phenochart 1.0.12 software (Akoya), a whole-slide contextual viewer was used for identification of regions of interest and acquisition of unmixed multispectral images.

### MALDI-TOF Mass Spectrometry Imaging (MSI) studies

#### Tissue Processing for MSI

Fresh human tonsils were snap frozen in liquid nitrogen following excision as described above. Tissue sections of 10 μm thickness were prepared using an HM525 NX cryostat (Thermo Scientific, Waltham, MA) operated at −20 °C, and thaw-mounted onto the conductive side of indium tin oxide (ITO)-coated slides. Slides were placed in a vacuum desiccator at −10 mmHg for at least 15 minutes prior to matrix deposition. Matrix deposition was performed with a TM-Sprayer (HTX Technologies, Chapel Hill, NC). For positive mode mass spectrometry imaging (MSI) experiments, 2,5-dihydroxybenzoic acid (DHB; Alfa Aesar, Ward Hill, MA) was sprayed onto the glass slide after tissue mounting at a concentration of 30 mg mL^-1^ with 12 passes/cycles. For negative mode MSI, 1,5-diaminonaphthalene (DAN; Sigma-Aldrich, Burlington, MA) was sprayed onto the glass slide after tissue mounting at a concentration of 5 mg mL^-1^ with 8 passes/cycles. Universal sprayer settings were as follows: nebulization gas pressure = 10 psi, track spacing = 3 mm, velocity = 1200 mm min^-1^, CC pattern.

#### Data acquisition and analysis

Data acquisition was carried out using a rapifleX MALDI TissueTyper tandem time-of-flight (TOF/TOF) mass spectrometer (Bruker Daltonics, Billerica, MA). Bare citrate gold nanospheres (nanoXact 60 nm, nanoComposix, San Diego, CA) were used for mass axis calibration in MSI experiments in a similar fashion to that described by Kolarova *et al.(19*) A mixture of phosphatidylcholines with known exact monoisotopic masses, created in-house, was used for mass axis calibration of lipid extract mass spectra. Acquired data were processed in FlexImaging 5.0 (Bruker Daltonics, Billerica, MA) and SCiLS Lab Pro 2020a (Bruker, Bremen, Germany). Putative lipid identifications were made using a combination of precursor ion *m/z* database searching (LMSD, lipidmaps.org) and *de novo* TOF/TOF product ion assignments. Mass spectra were adapted for size and clarity by regenerating figures using x- and y-coordinates exported from FlexAnalysis ver. 3.4 (Bruker Daltonics, Billerica, MA) and imported into OriginPro ver. 2021b (OriginLab Corporation, Northampton, MA). All mass spectra were normalized to the base peak. Peak picking was performed using the proprietary algorithm (SNAP) in FlexAnalysis ver. 3.4. Deisotoping was performed simultaneously within the same algorithm. Peak lists were generated for monoisotopic ions with signal-to-noise ratio greater than or equal to 2:1. A Venn diagram indicating peak list overlap between different sorted cell types was created using a free online tool from Ghent University Bioinformatics and Evolutionary Genomics laboratory (http://bioinformatics.psb.ugent.be/webtools/Venn).

#### Lymphocyte sorting and preparation for MALDI-TOF MS

Cells from two human tonsils were labeled and sorted as previously described (20). Briefly, PD1^hi^CD57^hi^ follicular helper CD4 T cells (T_FH_), CD19^hi^IgD^hi^ (naïve) and CD19^hi^IgD^low^CD20^hi^CD38^dim^ (enriched in germinal center-GC) B cells were sorted. The PD1^hi^CD57^hi^ tonsillar T_FH_ subset was chosen given its exclusive localization within the GC borders (20). Following sorting, cells were suspended in Mg^2+^ and Ca^2+^-free phosphate buffered saline (PBS, pH 7.4), flash frozen, and stored at −80 °C until further analysis.

#### Lipid extraction and analysis from tonsil tissue and sorted cells

Small pieces (~1 mg) of snap-frozen human tonsils were dissected from the intact specimen and homogenized in 400 μL of deionized water (11.2 MΩ, 4°C) followed by the addition of 800 μL of chloroform/methanol (1:2 v/v). Sorted B and T lymphocytes were similarly treated. The resulting mixture was vortexed (2500 rpm, 2 min), sonicated for 2 minutes and subsequently incubated on dry ice for 10 min. Next, 300 μL of chloroform/water (43:57 % v/v) was added and the final mixture was also vortexed (2500 rpm, 2 min) and sonicated for 2 min, then centrifuged (3000 rpm for 5 min at 4 °C) and the lower organic phase containing lipids dissolved in chloroform collected and kept at −80 °C. The remaining upper phase and precipitate were collected for lipids re-extraction by adding 500 μL of chloroform/methanol (1:2 v/v). This mixture was vortexed (2500 rpm, 2 min), sonicated (2 min) and incubated on dry ice for 10 min. Subsequently, it was centrifuged (3000 rpm for 5 min at 4 °C) and the lower phase collected and added to the first lipid extract. The total extract was stored at −80 °C prior to MS analysis. 10 mM stock solutions of DHB and 9-AA MALDI matrices in methanol were used for MALDI MS analysis of extracted lipids in positive and negative ion modes, respectively. 10 μL of each matrix solution was individually mixed with an equal volume of the lipid extracts (from either tonsil tissue or lymphocytes). The lipid/matrix solutions were vortexed for 3 min at 2500 rpm and 2 μL of the final solution was deposited on a MALDI AnchorChip target plate allowing the spots to co-crystallize at room temperature prior to MS analysis. 9-AA was used for negative mode extracts assay and gave better co-crystallization for spotted samples than previously observed with DAN.

### Statistics

Flow-cytometry and histocytometry generated data were analyzed using linear regression and the Wilcoxon and Kruskal-Wallis tests. Statistical analysis was performed with Prism software (version 6; GraphPad Software, La Jolla, California). Results were considered to be significant when p < 0.05. For MSI data, probabilistic latent semantic analysis (pLSA) was used to build a model with 5 components to differentiate the tonsillar follicle from the extrafollicular space. This was performed in SCiLS Lab Pro pLSA module. Box and whiskers plots for ions of interest with higher observed intensities in the segment defined as the follicle than in other tissue segments were also generated in SCiLS Lab Pro. Comparison of median intensity values was performed using a Kruskal-Wallis test.

## Results

### Assessing the tonsillar cellular composition

Flow cytometry-based, single cell analysis is the reference method for the phenotypic characterization of cells. We started our analysis by applying a multiparametric flow cytometry assay focusing on the immune cell types (T and B cells) that constitute the vast majority of non-adherent cells in human tonsils (Figure 1A). B cells (CD19^hi^) were further characterized by their expression of CD27, IgD, CD20 and CD38. An enrichment of the CD20^hi^CD38^dim^ subset (a phenotype compatible with GC localization (16)) was found specifically in the IgD^lo^CD27^dim/hi^ memory B cell compartment (Figure 1A). CD4 positive cells, which constitute the majority of CD3 T cells, present a subset expressing the unique phenotype PD-1^hi^CD57^hi/low^ (T_FH_) (20) (Figure 1A). A similar representation of memory B cell subsets was found while plasma cells (CD19^hi^CD27^hi^IgD^lo^CD20^dim^CD38^hi^) was the subset with the lowest frequency (Figure 1B). Cumulative data analysis showed a relatively high frequency of T_FH_ cells while the majority of CD8 T cells express a naïve (CD27^hi^CD45RO^lo^) phenotype (Figure 1B). A significant correlation was found between CD19^hi^CD27^hi/dim^IgD^lo^CD20^hi^CD38^dim^ (most likely resembling GC B cells) and i) T_FH_ subsets (CD4^hi^CD27^hi/lo^CD45RO^hi^PD-1^hi^CD57^hi^, CD4^hi^CD27^hi/lo^CD45RO^hi^PD-1^hi^CD57^lo^) ii) CD19^hi^CD27^hi/dim^IgD^lo^CD20^dim^CD38^lo^ (most likely representing naïve and memory non-GC B cells) and iii) plasma cells (Figure 1C). A mutual regulation between T_FH_ and GC B cells has been previously proposed (6). Our data indicate that such regulation takes place during the development of the tonsillar follicular/GC reactions.

**Figure 1.**
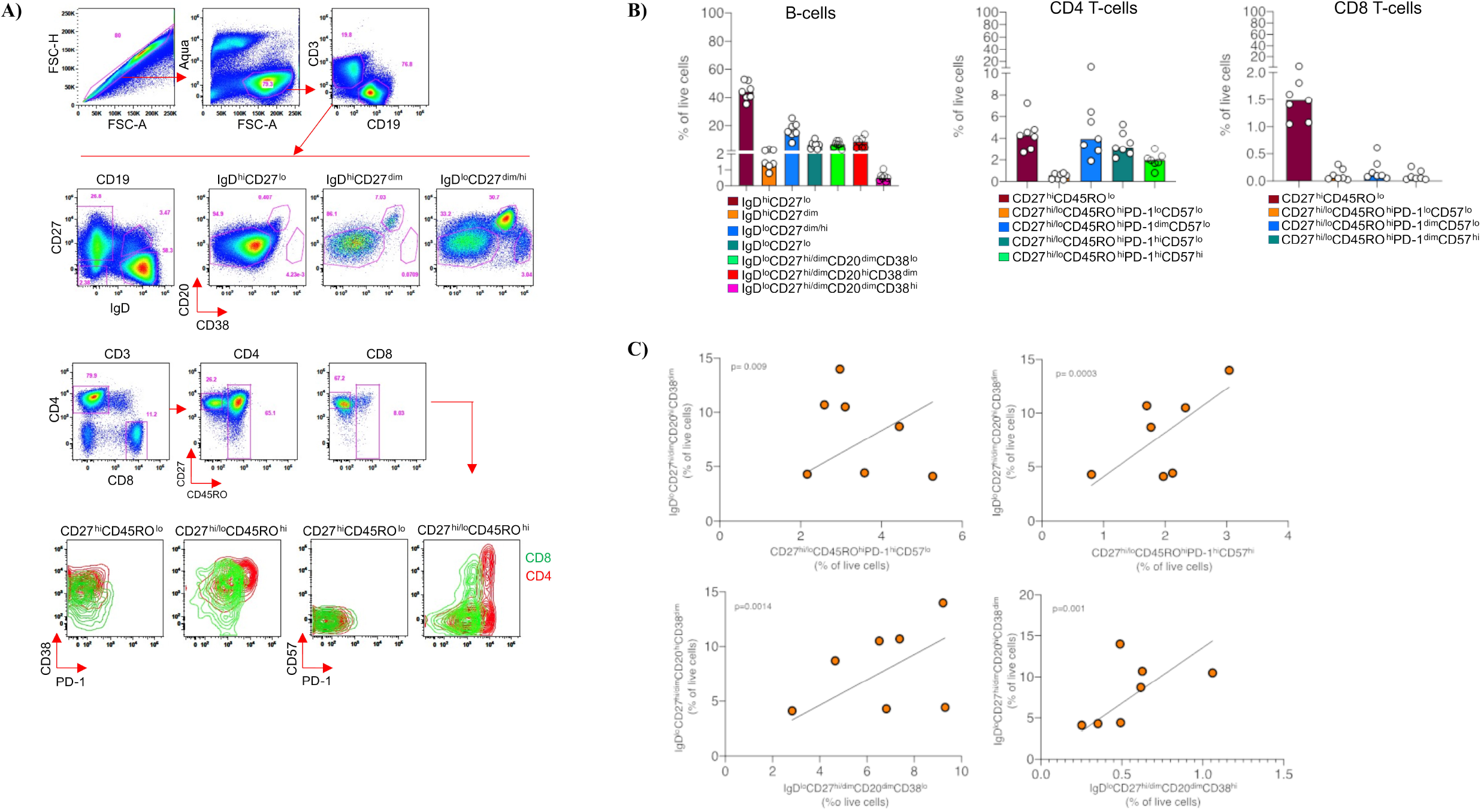
Assessment of the tonsillar cellular composition with multiparametric flow cytometry. **(A)** The gating scheme for the flow cytometry analysis of relevant cell subsets is shown (upper panel). The relative expression (intensity) of CD38, PD1 and CD57 in tonsillar CD4 (red) and CD8 (green) subsets is shown (lower panel). The overlay of corresponding 2D flow cytometry plots visualizes the different level of expression between the two T cell subsets with T_FH_ cells expressing the highest level of PD1. **(B)** Bar graphs showing accumulated data regarding the relative frequency of B, CD4 and CD8 subsets in tonsils (n=7) analyzed. (**C**) Significant correlation between GC B cells (CD19^hi^IgD^lo^CD27^hi/dim^CD20^hi^CD38^dim^) and i) TFH subsets (CD4^hi^CD27^hi/lo^CD45RO^hi^PD-1^hi^CD57^hi^ and CD4^hi^CD27^hi/lo^CD45RO^hi^PD-1^hi^CD57^l^°) (upper panel), ii) non GC memory B cells (CD19^hi^IgD^l^°CD27^hi/dim^CD20^dim^CD38^l^°) plasma cells (CD19^hi^CD27^hi/l^°IgD^l^°CD20^dim^CD38^hi^) (lower panel). Linear regression was used for the analysis.

### Skewed distribution of immune cell subsets and mediators of cell-to-cell communication among tonsillar areas

Despite the quantitative characterization of tonsillar cells by flow cytometry, this type of analysis does not provide any direct information regarding the localization of relevant cell subsets. Next, we sought to develop immunofluorescence multiplex imaging assays to address the localization of relevant cells and soluble mediators like the cytokine IL17 and chemokine CXCL13. Our confocal imaging assay allows for the simultaneous detection of i) B cell subsets, with respect to the expression of IgD, CD20 and Ki67, ii) CD4 T cell subsets, based on the expression of PD-1 and CD57, and iii) CD8 T cells and macrophages (CD68^hi^) (Figure 2A). The quantitative analysis of the obtained imaging data was carried out using the HistoCytometry pipeline (17, 21). The gating scheme for the identification of specific cell populations with respect to their area localization is shown in Supp. Fig. 1. Our analysis showed a highly skewed distribution of CD4, CD8 and B cell subsets across different follicular areas (Extrafollicular Area-EF, Mantle Zone-MZ and GC) (Figure 2B). Supporting the phenotypic characterization of the aforementioned B cell subsets (Figure 1A), CD20 expression was lower in the MZ (IgD positive area) and higher in the GC, particularly the LZ, in line with previous data (16, 20). Furthermore, the PD-1^hi^CD57^hi^ CD4 T_FH_ cells were found almost exclusively in the follicular area, specifically the GC (Figure 2B), in agreement with our recent data (20). On the other hand, CD8 T cells were highly excluded from the follicular areas while most CD68^hi^ macrophages were found in follicular areas (Figure 2B). A significant correlation was found between GC (CD20^hi^Ki67^hi^) B cells and PD-1^hi^CD57^hi^ CD4 T_FH_ cells (Figure 2B) in line with our flow cytometry data. The application of a second multiplex imaging assay revealed the localization of the FDC network within GCs, associated with an abundant expression of CXCL13 (Figure 2C), the main driving force for the trafficking of CD4 T cells and B cells towards the GC (22). On the other hand, IL17 expression was found almost exclusively in the Extrafollicular Area (Figure 2C), in line with our previously reported data related to the capacity of individual CD4 T cell subsets for IL17 secretion (23). Contrary to CD68^hi^ macrophages, CD14^hi^ monocytes were found to populate Extrafollicular Areas (Figure 2C). Therefore, in addition to the compartmentalization of immune cell types and soluble factors across the tonsillar areas, our data reveal the ‘exclusion’ of specific cells (*e.g*. CD8, CD14) from tonsillar GCs.

**Figure 2.**
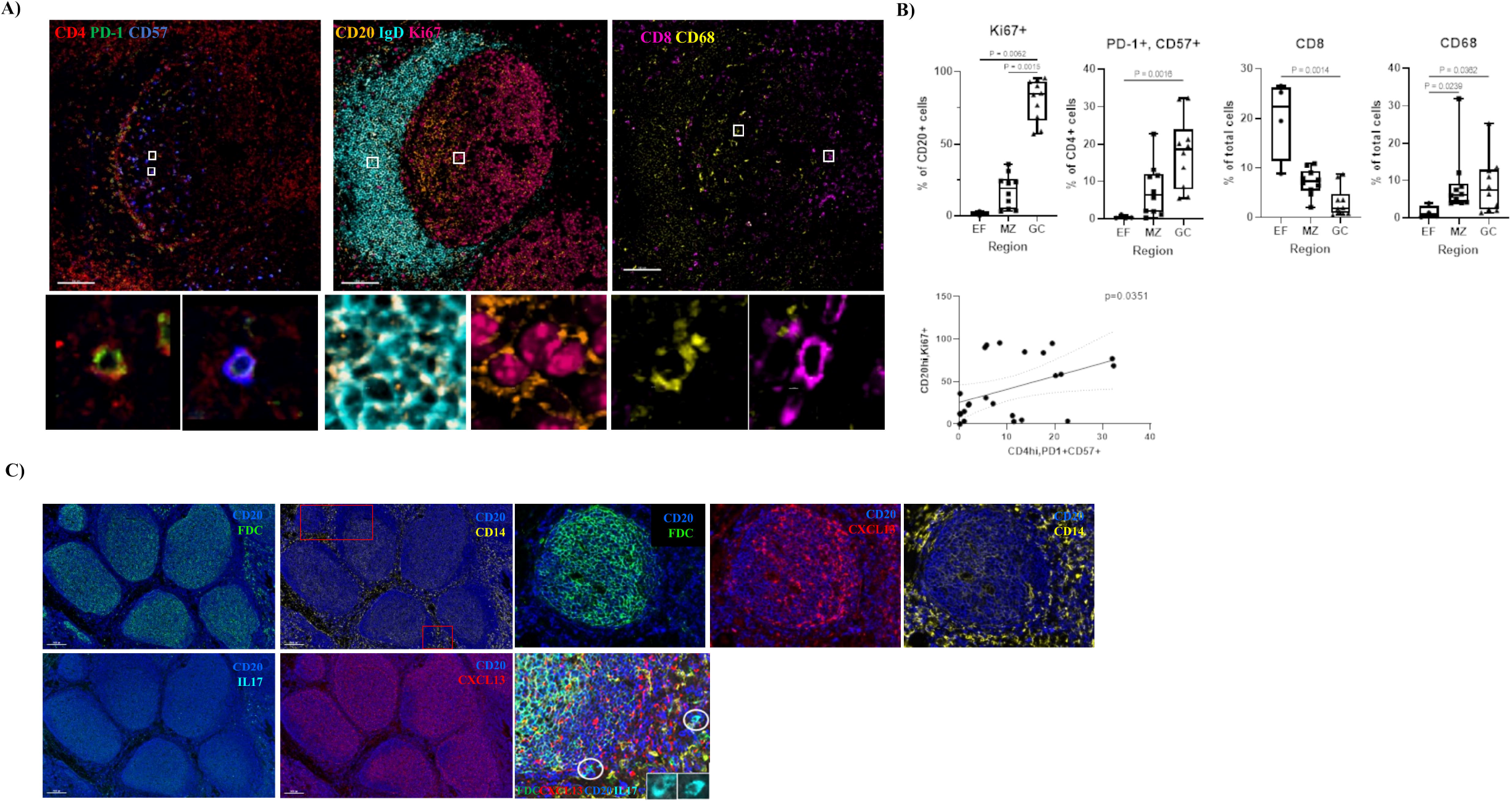
Compartmentalization of tonsillar cell types and soluble mediators revealed by multiplex imaging analysis. **(A)** Representative multiplex confocal images showing the distribution of B, CD4 and CD8 cell subsets in a follicular area and adjacent ‘T cell area’. Zoomed areas (white boxes) are also shown for the visualization of the expression of relevant biomarkers on individual cells. (**B**) Bar graphs showing the relative expression (as frequency of the corresponding parental population) of Ki67^+^ B cells and PD-1^+^CD57^+^ CD4 T cells as well as the frequency (of total cells) of CD8 and CD68 subsets in different tonsillar areas (EF=extrafollicular, MZ= mantle zone, GC= germinal center) (upper panel), Kruskal-Wallis statistic test used significant differences notated. The correlation between Ki67^+^ GC B cells and PD-1^+^CD57^+^ CD4 T_FH_ cells is shown (lower panel), Simple linear regression statistical test used. (**C**) Representative multispectral images showing the distribution of FDCs (green), CD14 (yellow), CXCL13 (red) and IL17 (cyan) with respect to the follicular areas (CD20, blue) in one tonsil. Inserts are showing the expression of IL17 in zoomed areas (white circles).

### Compartmentalized localization of tonsillar lipids

Then, we sought to investigate the *in situ* distribution/localization of lipids, which represent critical components for the structure of cellular membranes, intracellular signaling, and communication between cells (chemo-attractants) (24). The general scheme of the experimental pipeline for lipid imaging using MSI and subsequent ion identification by MS/MS is shown in Supp. Fig. 2A. First, MSI was performed to identify regions and compounds of interest using a single stage mass spectrum. Multiple ion images (MII) were created manually to find a sufficient number of ions to fill the entire imaged tissue area and show the tissue morphology. Individual extracted ion images (EII) were used to identify pixels with high intensity ions of interest. An on-tissue, space-directed MS/MS approach was used to reduce combined fragmentation of isobars (ions with different chemical structures but the same nominal mass) which may be found overlapping in the average mass spectrum but are distributed differently throughout the tissue. Briefly, collisional activation and subsequent spontaneous decomposition was performed in the collision region of the TOF/TOF after focusing the laser in a single 20 μm beam on the region where the compound of interest was shown to reside in the sample. Precursor ions were given putative structures based on their characteristic product ions (*e.g*. polar headgroup product ions for glycerophospholipids). We started our investigation by analyzing lipids with MSI in positive mode, using DHB matrix, data acquisition at 20 μm lateral resolution (Supp. Fig. 2B), and data normalization using root mean square (RMS) correction (Supp. Fig. 2C). The average single stage mass spectrum across the entire imaged tissue (inset) and a zoomed region from *m/z* 700 to 750 to indicate the ions used for further analysis, are shown (Figure 3A). In total, 74 common monoisotopic *m/z* peaks were detected when the mass spectrum was averaged across the entire tissue and compared between different tonsillar specimens (N=3). Twenty of these peaks were found to be localized to the tissue segment designated as the follicular area. These peaks, identified as lipids, did not always display a similar tissue distribution but in all cases were more abundant in the follicular area than in other tissue regions (Supp. Fig. 3). An exhaustive characterization of all detected ions, with treatment of each spectrum separately, may be necessary for discovery of novel biological phenomena in the future. Most of the abundant peaks in the spectrum correspond to protonated ([M+H]^+^) phosphatidylcholines (PCs), with less frequent peaks corresponding to sodiated ([M+Na]^+^) diacylglycerols (DAGs), protonated sphingomyelins (SMs), and lyso-lipids (Supp. Fig. 3). SMs are observed with the same ionization characteristics as PCs due to their identical phosphocholine head group; therefore, they are not easily differentiated in positive mode MS. Using the nitrogen rule (25), inferences were made regarding the identities of these ions (*i.e*. odd mass ions were inferred to be primarily from SM monoisotopic ions and even mass ions were inferred to be primarily from PC monoisotopic ions). Collisional activation and dissociation of these ions suggests that some include isobaric isotopic peaks from other species. We focused on three ions (*m/z* 703, 725, and 728), based on their distribution profile, for further analysis. The localization of these ions across the entire tissue resembles tonsillar epithelium or connective tissue, extrafollicular and follicular areas, respectively (Figure 3A), and they were putatively identified as SM 34:1, SM 36:4, and PC 32:3 using precursor ion mass and product ion spectra from the same instrument as was used to acquire images. Their tandem mass spectra (LIFT MS/MS) and respective zoomed images to better visualize the tissue micro-anatomy are shown in Figure 3B. The magenta boxes indicate the ion channel at *m/z* 184 (phosphocholine ion), identifying these ions as either PCs or SMs. The corresponding structure of these three lipids is also shown (Figure 3C). The structure identifications are putative as there could be several isomers or isobars isolated and fragmented in the tandem mass spectra. The major lipid classes in the tandem mass spectra are defined by the polar lipid headgroup product ions indicated.

**Figure 3.**
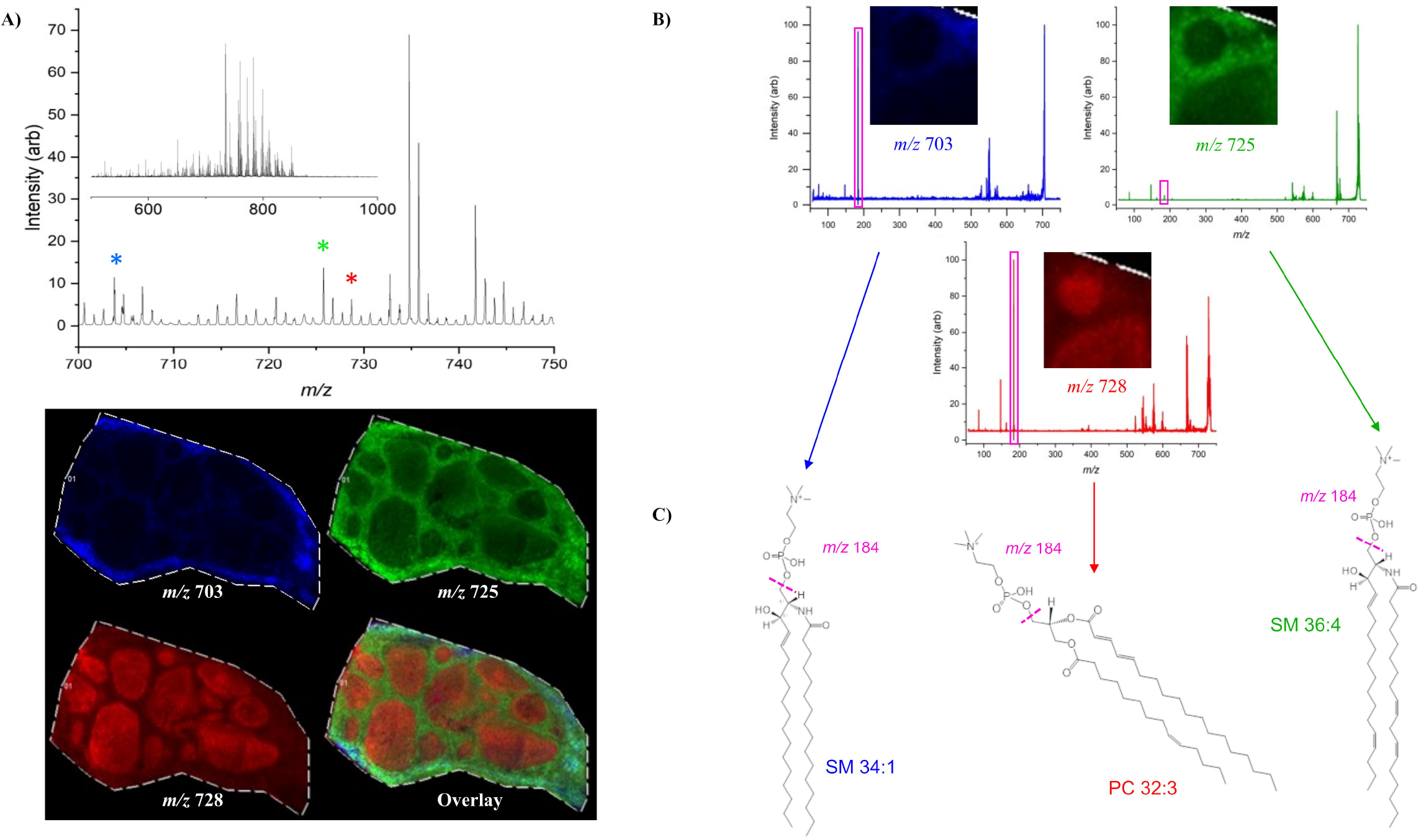
Average positive ion mode single stage TOF mass spectrum showing lipids detected over the mass range *m/z* 500-1000 **(A, top inset)** using DHB as MALDI matrix. Lipid ions used to create images **(A, bottom)** are indicated in the region *m/z* 700-750 with asterisks corresponding to the colors used to generate images **(A, top)**. Tandem mass spectra for each of the three lipids **(B)** were acquired to determine lipid class. Magenta boxes indicate the characteristic phosphocholine ion at *m/z* 184 determining these ions as either SMs or PCs. Zoom regions of the images around two tonsillar follicles are shown for clarity. Putative structure identifications using precursor ion mass and headgroup product ion mass are shown for each of the three ions **(C)**. Odd mass precursor ions were inferred to be from SMs and even mass precursor ions were inferred to be from PCs. Lipid isomers may be present in the sample and acyl chain double bond positions indicated in the figure are not determinable using the positive ion mode method applied.

Next, negative mode MSI was performed using DAN matrix and imaging at 20 μm lateral resolution (Supp. Fig. 4A). In total, 80 common monoisotopic *m/z* peaks were detected when the mass spectrum was averaged across the entire tissue and compared between patient specimens (N=3). Twenty-four of these peaks were found to be localized to the tissue segment designated as the follicular area (Supp. Fig. 4B). We further focused our analysis on seven ions, putatively identified as lyso-SM d18:0, PC O 20:4, LPI 18:0, SM 34:1, SM 36:1, SM 42:2, and PI 34:1 (Figure 4A, upper panel). Their localization across the imaged tissue is shown (Figure 4A, lower panel). The tandem mass spectra (LIFT MS/MS) for four of the ions imaged (Figure 4B), and their basic structure elucidation scheme (Figure 4C) are shown too. The magenta boxes (Figure 4B) indicate the ion channels corresponding to the glycerophospholipid head group ions used to identify the lipids. Figure 4C shows representative lipid structures for each of the precursor ions in Figure 4B. These are putative identifications because there could be several lipid isomers or isobaric compounds isolated and fragmented in the tandem MS experiments. However, the major lipid class was positively identified by the headgroup product ion. Collectively, our data demonstrate that tonsillar areas are characterized by distinct positioning of many lipid molecules, possibly contributing to the structural and functional compartmentalization of this lymphoid organ.

**Figure 4.**
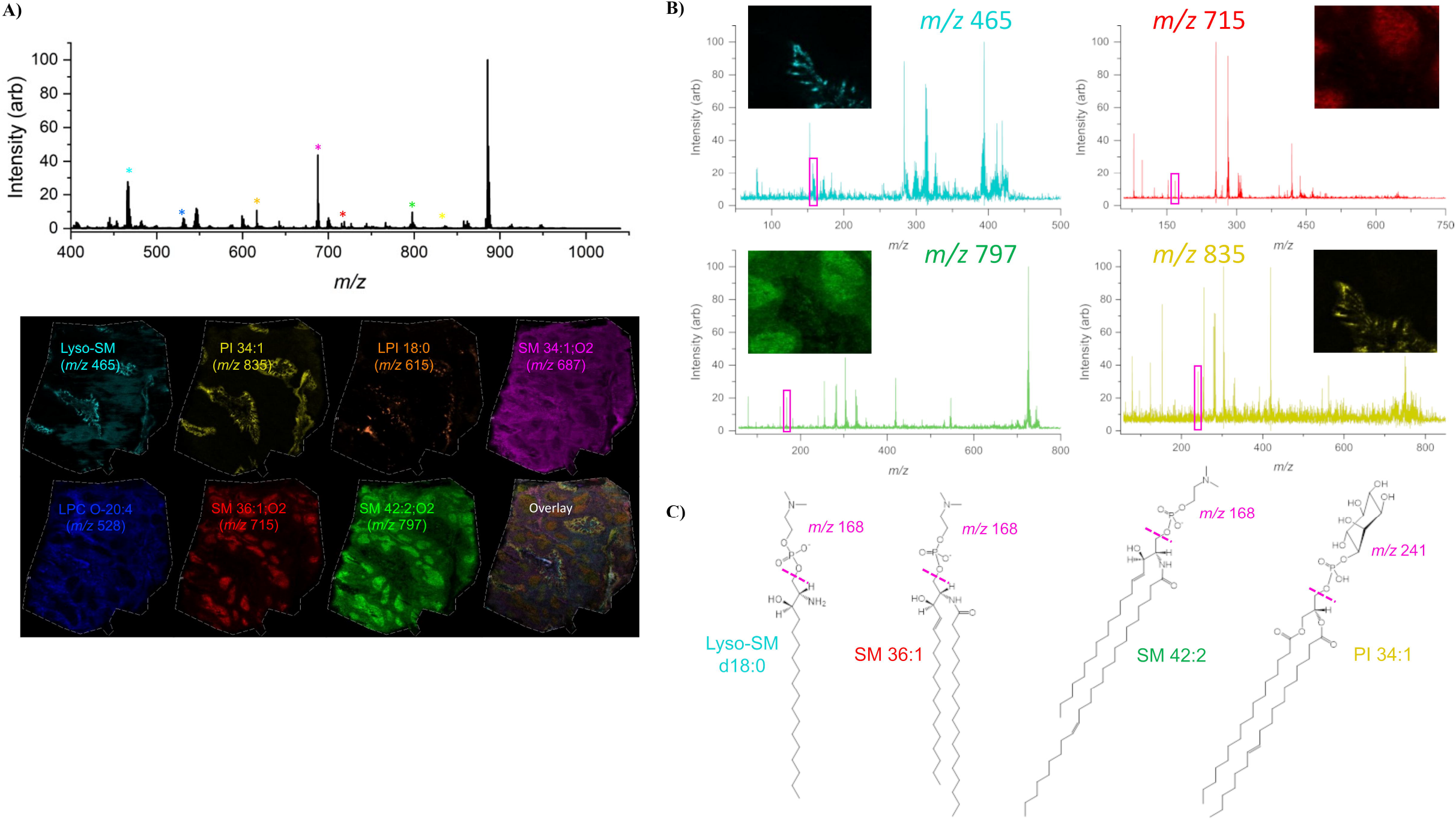
Average negative ion mode single stage mass spectrum in the mass range *m/z* 400-1050 using DAN as MALDI matrix **(A, top)**. Ions used to generate images in the bottom panel are indicated with asterisks in their corresponding colors in the mass spectrum. Extracted ion images (EII) for each of the lipid ions indicated are shown **(A, bottom)** along with an overlay of the seven ions chosen to create the histological image. **(B)** Tandem mass spectra for four of the ions chosen to create the tissue image are shown with magenta boxes indicating the polar lipid headgroup product ion used to identify the lipid class. In the region *m/z* ~200-315, fatty acyl carboxylate anions can be seen in the negative ion mode tandem mass spectrum. While the posititions of acyl chain double bonds cannot be determined using this method, SMs can be differentiated from PCs based on their lack of acyl chain product ions in contrast to positive ion mode where they cannot be differentiated. **(C)** Basic structure elucidation schemes with representative structures for the ions imaged are shown. The phosphocholine product ions from PCs and SMs in negative ion mode always lose a methyl group, indicated by the change in mass from *m/z* 184 to *m/z* 168. The phosphoinositol headgroup mass in negative ion mode is *m/z* 241, along with other small indicative phosphoinositol fragments.

### Computational analysis of lipid compartmentalization

Computational tissue segmentation, for both positive and negative ion modes, was performed using the SCiLS Lab Pro Segmentation module. Two different *m/z* ranges were used to automatically segment the tissue in negative ion mode. Use of the 300-1100 *m/z* range resulted in poor tissue segmentation while the 400-1100 *m/z* range provided a segmentation resembling the tissue micro-anatomy observed by the corresponding imaging of tissue using a nuclear marker (Supp. Fig. 5A). It is likely that extending the mass range below *m/z* 400 included matrix ions in the segmentation, a confounding factor for the segmentation algorithm. Automatically generated tissue segments were further assigned as tissue regions in the software. The three tissue segments assigned to the tonsillar follicle, extrafollicular area, and tonsillar crypt were used for further processing and statistics. Receiver operating characteristic (ROC) curves were used for the validation of a given ion to discriminate between the tonsillar segments (Figure 5A). An ion specific for the crypt and an ion that uniformly distributed across the tissue gave AUC values of 0.207 and 0.580, respectively, while a follicle-specific ion gave a value of 0.957, associated with a distinct distribution pattern primarily within the follicle (Figure 5A). An AUC value of 1.000 in this analysis is considered ideally specific and sensitive to distinguish one segment from the other. A probabilistic latent semantic analysis (pLSA) was performed to identify lipids that could contribute to the differential imaging profiles across follicular subareas. Similar to a principal components analysis, a pLSA creates “components” or vectors of the data set with discriminatory power and plots data groups on axes created from those components. Discriminatory power of each individual ion and its contribution to the component can be extracted and used for biomarker discovery, if needed. A bimodal distribution of ions was observed in the pLSA (Figure 5B, upper panel), indicating a differential expression of lipids among follicular areas, likely GCs and MZs in this case (Figure 5B, lower panel). Representative tissue segmentation images are shown in both positive and negative ion modes for a visual reference of the segmentation performance (Figure 5C, left panels). The tissue segmentation maps were generated with the identified regions arbitrarily given different colors for contrast. Regenerated images using the negative ion mode pLSA data points from two of the five model-generated “components” in the region defined as the follicle are also shown in Figure 5C (right panels). These two components indicate that the follicle could potentially be segmented further into areas which may correspond to the GC and MZ. Multi-modal imaging of the same tissue section would be useful to test this hypothesis and further validate the data analysis method. Tissue segmentation maps were generated automatically in SCiLS Lab Pro using all the features (*m/z* channels) detected in the imaging run, pixel-by-pixel. Pixels with similar features and relative abundances of those features were grouped together as segments. Some segments were then grouped together manually to represent the known tonsillar tissue architecture.

**Figure 5.**
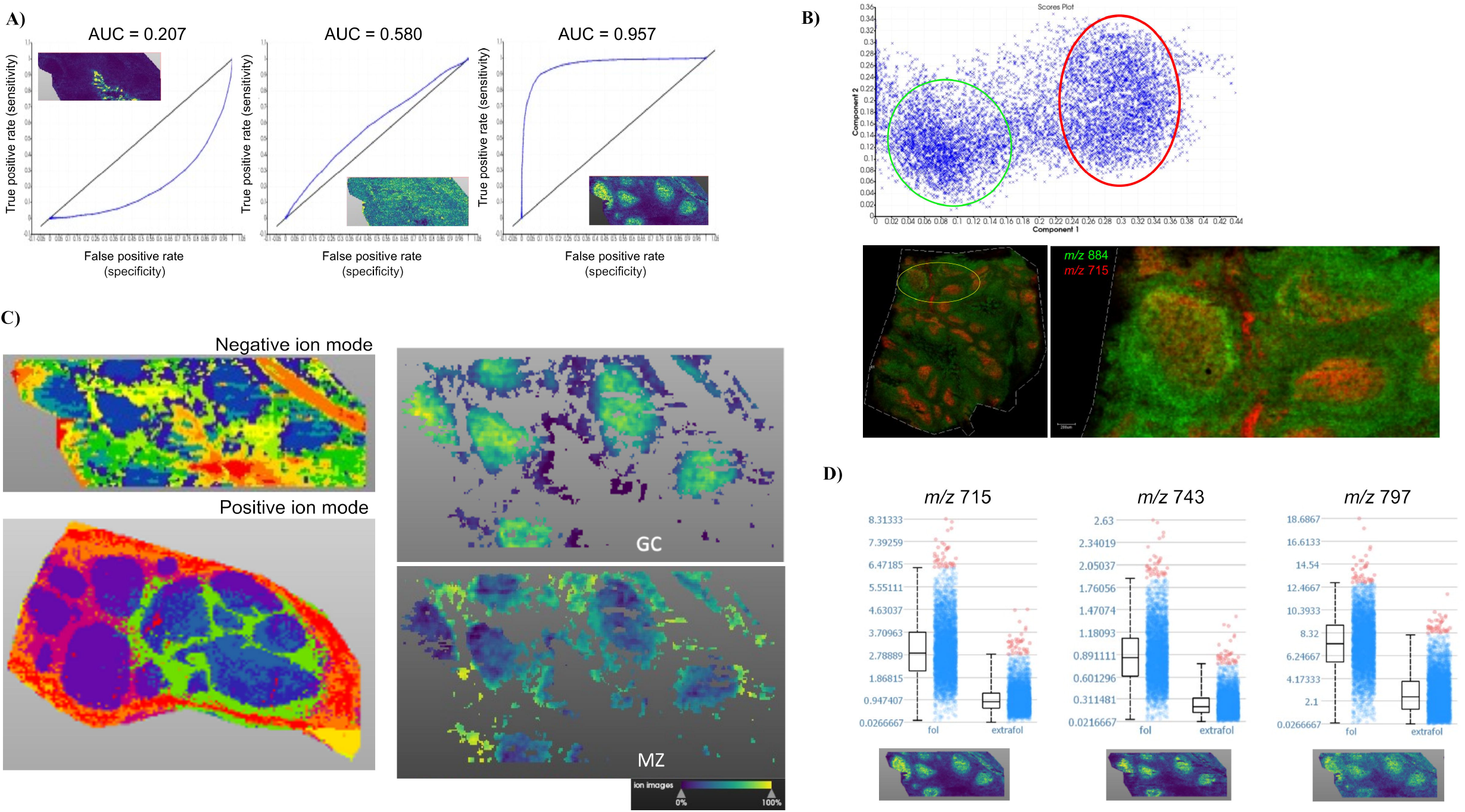
**(A)** Receiver operating characteristic (ROC) curves for three ions in negative ion mode when comparing the specificity of each ion for the follicular area versus the extrafollicular area. Area under the curve (AUC) for an ion specific to the tonsillar crypt/epithelium was 0.207, indicating it is not specific to the follicle. AUC for an ion found relatively evenly distributed across the tissue was 0.580, indicating that it is found in the follicle but not specific to that area versus the extrafollicular area. AUC for an ion specific to the follicle was 0.957, indicating its ability to differentiate the follicle from the extrafollicular area. Images of the three ions used to generate the ROC curves are shown as insets. **(B)** Probabilistic latent semantic analysis (pLSA) using a model with five components showing a bimodal distribution of data points in the region defined as the follicle (top). Circles were drawn around the data points for emphasis. Two ions, *m/z* 715 and *m/z* 884, are representative of two regions of the follicle computationally derived and shown for analysis clarification (bottom). **(C)** Computational segmentation of tonsillar tissue in negative and positive ion modes using all features in each individual mass spectrum from the imaging experiments (left panels) and computational regeneration of the data from the pLSA indicating the region defined as the follicle could be further divided into two distinct regions (right panels) inferred to be the germinal center (GC) and the surrounding mantle zone (MZ). **(D)** Three ions (*m/z* 715, 743, and 797) found to be specific to the follicle versus the extrafollicular space, but demonstrating unique tissue localization images (bottom) indicating that these lipids are compartmentalized slightly differently in the tissue. Box and whiskers plots show that in all cases the median intensity values of these ions are higher (p < 0.002 for all three ions) in the follicle than in the extrafollicular space.

We further explored the distribution of three ions (*m/z* 715, 743, and 797) preferentially localized to the follicle relative to the extrafollicular area. Their relative quantification revealed a significantly (p < 0.002 for all 3 ions) higher median presence in the follicle than in the Extrafollicular Area, in two different patient specimens (Figure 5D and Supp. Fig. 5B), which is a computational result consistent with what is observed with the eyes in the corresponding images (Figure 5D and Supp. Fig. 5B). Therefore, the shown computational analysis represents a robust pipeline for the relative quantitative analysis of a given lipid with respect to its tissue localization.

### Comparative lipidomic profiling of tonsillar cell subsets

Access to fresh tonsillar tissues allowed for a comparative lipidomic profiling approach, utilizing specific, flow cytometry sorted CD4^+^ and B cell subsets for MALDI-TOF mass spectrometry and MSI of sections from the same tissue (Supp. Fig. 5C). Tonsillar cells were sorted based on known receptors for follicular helper CD4 T cells (T_FH_) (CD4+CD27+CD45RO+PD1^hi^CD57^hi^, a GC enriched T_FH_ subset (20)) and B cells (naïve-CD19^hi^IgD^hi^, GC enriched-CD19^hi^CD20^hi^CD38^dim^). First, the mass spectra (positive and negative ion mode) for unfractionated tonsillar extracts were generated by MALDI-TOF (Figure 6A). A comparison of the negative ion mode single stage mass spectra acquired for the entire tissue section and the corresponding unfractionated tonsillar extract are shown (Figure 6B). Although more features were observed in the spectrum for extracted tissue compared to the imaging profile, the major features remained the same (*e.g*. the base peak in all average negative ion mode spectra was *m/z* 885, the “canonical” phosphatidylinositol (PI 38:4) ion). Our cell-based lipidomic profiling represents the relative abundance of a given lipid for the entire tissue while its MSI analysis only for the imaged plane which may not always represent the distribution of a lipid across the whole tissue. Positive ion mode spectra comparisons between MS images and cellular extracts were also similar in both features detected and relative abundances; negative ion mode spectra were chosen for display in Figure 6B due to reduced dominance of the PC ion series. These data serve to further confirm MSI results with respect to compounds detected and their relative abundances in the tissue. The averaged mass spectra for three sorted lymphocyte cell subsets and respective images of three ions (*m/z* 715, *m/z* 747 and *m/z* 882) found in those cells and localized to regions where they are known to reside are shown in Figure 6C. Overlays are also shown to highlight the unique tissue distribution of these ions with respect to the others (Figure 6C). In total, 57, 51, and 48 monoisotopic ions were detected in negative ion mode uniquely in the GC CD4 T cells, naïve B cells, and GC B cells, respectively (Figure 6C). Since we were only comparing these three cell types to each other, this does not necessarily suggest that all these ions are unique to one human cell type. However, it is possible that a subset of these ions could be used for phenotypic characterization complementary to other molecular analysis methods. Our data demonstrate that the complementary application, when access to tissue-derived cells is possible, of mass spectrometry-based cell phenotyping and MSI methodologies could provide meaningful information regarding the lipid spatial localization and cellular origin with high specificity based on the markers used for sorting.

**Figure 6.**
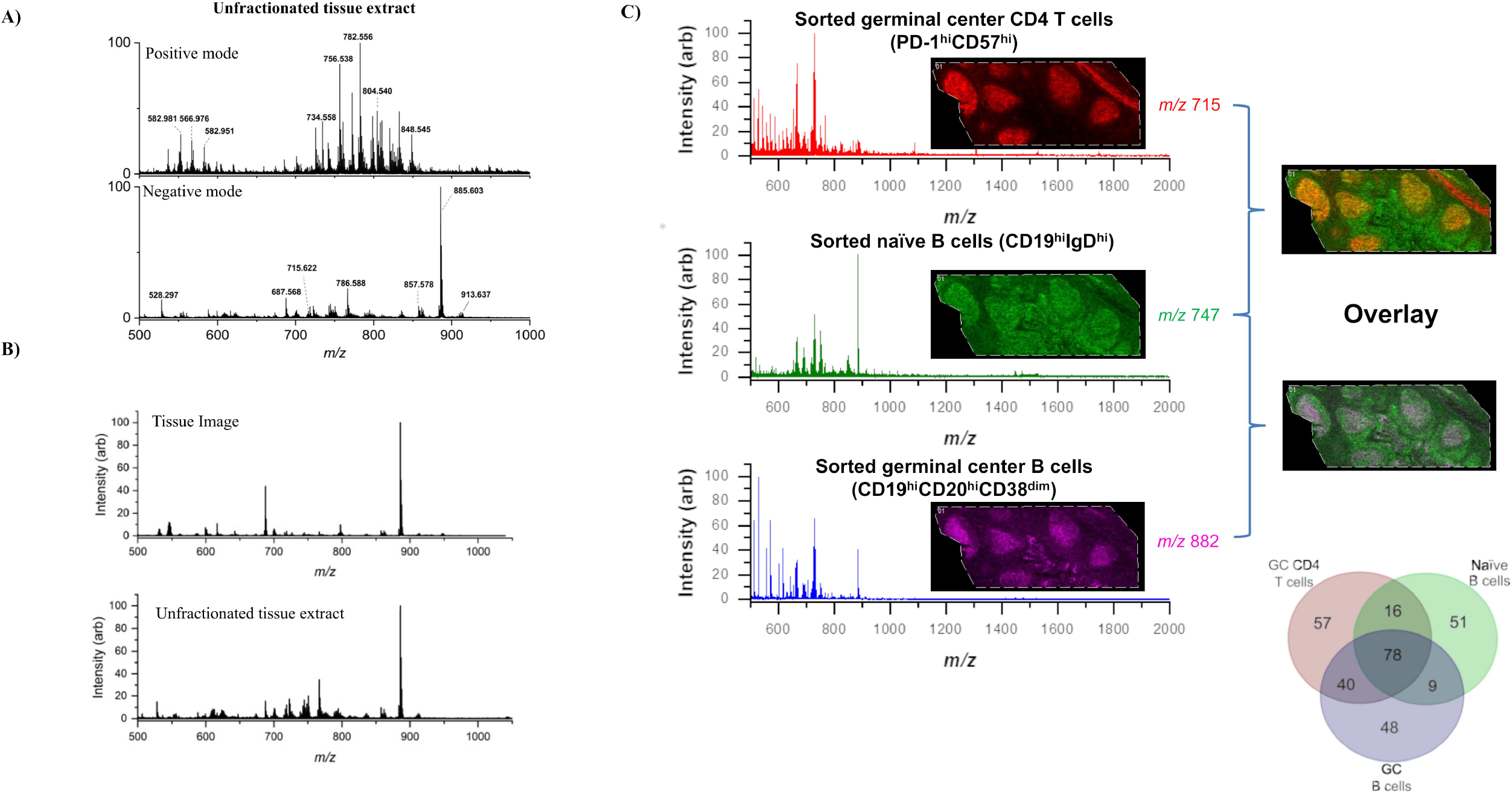
**(A)** Broadband single stage mass spectra in positive (top) and negative (bottom) ion modes for unfractionated tonsil tissue extract using DHB and 9-AA as matrices in positive and negative ion modes, respectively. Small pieces of tissue were dissected from a frozen tonsil and extracted using a total lipid extraction method. **(B)** Average broadband single stage mass spectra acquired in negative ion mode from a tissue image (top) versus an unfractionated tissue extract (bottom). While more features were detected in the tissue extract, and some with slightly different relative abundances, the major features and their relative abundances in the mass spectra were strikingly similar. **(C)** Mass spectra shown are broadband negative ion mode spectra from the sorted tonsillar cell subsets: GC CD4 T cells, naïve B cells, and GC B cells. Each spectrum is obviously different, indicating that these spectra could be used as molecular barcodes to identify a particular cell phenotype. Three ions (*m/z* 715, 747, and 882) with different tissue distribution in the images and found in different relative abundances in each cell phenotype are shown to indicate that these ions may be specific to those cell phenotypes in comparison to the others (this cannot be extrapolated to comparisons of cell phenotypes not investigated in this study). A Venn diagram showing the monoisotopic lipid ions detected in each cell type and their uniqueness and overlap is presented for emphasis that the lipid mass spectra from each cell type are distinguishable from one another.

## Discussion

Lymphoid follicles are the micro-anatomical sites where B-cell maturation and the development of antigen-specific B cell responses occur. The development of these responses is a multi-step process wherein a wide range of innate and adaptive immune cell subsets function in a coordinated mode. Within the GCs, the quality of the T_FH_ “help” received by GC B cells ultimately determines the quality of B cell responses (26). The different phases of this process are mediated by interactions between immune and non-immune cell subsets and take place in distinct lymphoid tissue areas. This compartmentalization of immune interactions could facilitate i) the ordered interaction between different immune cell subsets as well as between immune cells and other tissue determinants (*e.g*. stromal cells FRC, FDCs), and ii) the creation of multi-step checkpoints that could prevent unwanted immune reactions (*e.g*. autoimmunity, presence of cytotoxic cells like CD8 and NKs in GCs). The relatively easy access, the preservation of distinct anatomical areas, and the presence of numerous mature follicles make tonsil an ideal tissue for the understanding of SLO immune organization. In line with our previous data (20), we found a high representation of tonsillar T_FH_ subsets (PD-1^hi^CD57^lo^, PD-1^hi^CD57^hi^), while the majority of CD8 T cells express a naïve phenotype, suggesting a limited ‘Th1’ type immune reactivity in this tissue environment. A balanced representation among different B cell subsets was found, while a significant correlation between GC B cells and T_FH_ subsets, especially the CD57^hi^ ones, or plasma cells was observed. Therefore, the cellular composition determined by flow cytometry points to a coordinated development, possibly supported by mutual regulation (6) of major tonsillar GC cell players and could set the stage for similar comparisons in lymph nodes from diseased subjects. Given the use of tissue-derived cells, flow cytometry provides a relatively robust estimation of cell subset frequencies across the tissue, a measurement that could be used as a reference for the interpretation of quantitative data from imaging analysis when tissue-derived cells are available.

We applied a quantitative multiplex imaging pipeline for the analysis of the spatial organization of major cell subsets in tonsillar areas. A skewed distribution of several cell subsets analyzed was observed among the main tonsillar areas (Extrafollicular, MZ, GC). Our data confirmed the relative absence of CD8 T cells from the GC area, a profile compatible with ‘immune privileged’ sanctuary areas, where the uncontrolled presence of cellular sources of killing mediators like FasL could be detrimental for the local immune cell subsets (T_FH_, GC B cells) that overexpress Fas (27). Similar to flow cytometry data, a significant correlation between T_FH_ and GC B cells was found. Contrary to CD14^hi^ monocytes, a skewed representation of CD68 within the GC and MZ was found, marking the so-called tingible-body macrophages (28). Besides the compartmentalized positioning of cell types, we found a differential expression of secreted mediators like CXCL13 and IL17. The skewed positioning of CXCL13 is compatible with its function (trafficking of T_FH_ and B cells through CXCR5). The absence of IL17 from the follicular area, particularly GC, is in line with our previous observation that T_FH_ subsets do not secrete this cytokine (23). We assessed the expression level per cell for CD20, a widely used B cell marker, between different follicular areas. We found a diminished expression in the MZ (IgD^hi^ B cells) while the highest expression was found in the LZ. Therefore, the quantitative imaging data are complementary to those generated by flow cytometry and can facilitate the assignment of specific tissue positioning to B-cell phenotypes identified by flow cytometry. This is in line with the recently described positioning of CD57^lo^ and CD57^hi^ T_FH_ cells across the follicular areas (20). Therefore, the combinational use of flow cytometry and multiplex imaging can significantly increase our capabilities for understanding the tissue immune system organization.

Due to their fluorescent labeling paradigm, flow cytometry and confocal tissue imaging can provide only ‘targeted’ data with respect to the molecules/markers under investigation. In addition, these methodologies cannot provide any information for molecules without a known specific probe (*e.g*. lipids and metabolites). Lipids are abundant molecules that can mediate a wide range of functions including formation of cellular membranes, signal transduction, and chemo-attraction (*e.g*. lymphoblast chemo-attraction by human plasma via lysophosphatidylcholine (29)). We applied a mass spectrometry-based imaging platform to access the distribution of several lipids across tonsillar areas. To this end, we performed an untargeted screening MSI protocol using both positive and negative ion modes. Advantages of negative ion mode, when analyzing glycerophospholipids under the conditions described, are i) the reduced spectral complexity due to ionic adducts and ii) the reduced dominance of PCs in the mass spectrum as compared to positive ion mode. One advantage of positive ion mode lipid imaging is the detection of metal ion adducts from compounds otherwise unionizable by MALDI (*e.g*. DAGs and TAGs). A limitation to both modes on the instrument used for these analyses is the lower mass resolving power than is required for isotopic fine structure analysis and separation of isobaric species. Often this is required for positive identification of compounds in complex mixtures with no separation prior to the mass analyzer. Since MSI is performed *in situ*, the only currently available instruments that can separate isobaric ions prior to the mass analyzer are hybrid ion mobility spectrometer-mass spectrometers. Another commonly employed strategy for positive identification of compounds is to use a higher mass resolving power (but lower throughput) mass spectrometer for MSI experiments; however, this strategy often does not overcome uncertainty associated with isomers (compounds with the same chemical formula and exact mass).

Previous work on lymphoid tissue lipidomics/metabolomics is sparse. However, human formalin-fixed paraffin embedded (FFPE) tonsils sections have been mapped using time-of-flight secondary ion mass spectrometry (TOF-SIMS), a high lateral resolution MS imaging technique (30). 189 metabolites in tonsils were found and some of them were associated with T and B lymphocytes in tonsils by using isotope-labeled antibodies such as anti-CD4 and anti-CD20, respectively. The major limitations to TOF-SIMS are its low upper limit of molecular weight ions that can be detected using the commonly employed methods, and extensive in-source fragmentation of compounds by the particle beam; therefore, only lipid fragments and other small metabolites were detected in the aforementioned study. MALDI-TOF MS has a much higher high mass cut-off in the hundreds of kilodaltons and much less in-source fragmentation, allowing detection and relative quantification of a much more diverse repertoire of compounds. This leads to fewer inferences as to the origins of fragment ions detected. In contrast to the previous study, Yagnik *et al*. (31) proposed a multiplexed immunohistochemistry (IHC) method based on MALDI imaging to detect immune cells such as helper T cells, cytotoxic T cells, memory T cells and B cells in FFPE tonsil tissue by using a tissue labeling approach with photocleavable isotope-labeled fluorescent antibodies, allowing liberation and detection of mass tags once the laser irradiates the sample. Denti *et al*. also performed a lipid signature study of T lymphocytes found in FFPE human tonsil tissue by using MALDI-MSI and IHC (32). They found that glycerophospholipids such as phosphatidic acids (*m/z* 701.5, PA 18:0/18:1; *m/z* 723.4, PA 20:4/18:0; *m/z* 725.4, PA 18:0/18:3) and phosphatidylethanolamine (*m/z* 702.6, PE 40:2) were highly abundant in T-cell-rich regions identified by CD3, CD4 and CD8 antibodies but lower in epithelial regions.

We found a compartmentalized distribution of several lipids analyzed. Using three ions, *m/z* 703, 725 and 728, representing SM 34:1, SM 36:4, and PC 32:3, respectively, we were able to computationally segment the tonsillar epithelium/crypts, extrafollicular, and follicular areas in positive ion mode. In negative ion mode, we were able to produce similar results with enrichment for non-PC lipids and lyso-lipids. There were 24 ions and 20 ions detected in positive and negative ion modes, respectively, with a more abundant distribution in the follicular areas than in the extrafollicular areas. Unsurprisingly, phosphatidylcholines (PCs, ~ *m/z* 700-800), major constituents of the total membrane mass (33) and essential nutrients required to maintain lymphoid cells in culture (34), were found to be the most abundant lipid ions detected in positive mode MSI experiments. In fact, PCs are usually so dominant in positive mode MALDI-TOF mass spectra that many other cationizable lipids may be suppressed. In negative ion mode MALDI, both PCs and SMs lose a methyl (-CH_3_) group from the quaternary amine choline moiety, rendering it neutral, and the single negative charge resides on the phosphate moiety. Negative mode tandem mass spectrometry (MS/MS) is able to discriminate PCs from SMs by presence/absence of fatty acyl carboxylate anions in the tandem mass spectrum (*i.e*. PCs are able to liberate ionizable acyl groups through collisional activation while SMs cannot). Both PCs and SMs are identifiable by a phosphocholine-minus-methyl fragment (*m/z* 168), a phosphite ion (*m/z* 79), and a phosphonate ion (*m/z* 97) in the tandem mass spectrum. Unlike PCs, increased membrane PIs impart net negative charge to the cellular membrane which affects membrane fluidity, shape, and the types of other non-PI components found in PI-rich membrane loci (35). They are observed in negative ion mode as singly deprotonated ions with high sensitivity. This is an advantage to using negative ion mode for PI analysis since PIs have no proton acceptor groups; they are detected in positive mode, if at all, as cation adducts which requires further spectral interpretation. PIs are also the precursors of phosphoinositides, kinase-phosphorylated PIs that are important mediators of many G protein-coupled receptor (GPCR) reactions in the cytosol (36). A PI at *m/z* 882 (likely an isotopic peak with less isobaric interference than its respective monoisotopic peak), confirmed by MS/MS and found to be selectively expressed in GCs, was also found in sorted GC B cells. Furthermore, the pattern was not homogeneous across the follicular area, indicating that this lipid could be differentially expressed in the LZ and DZ.

Co-registration experiments using antibodies against protein biomarkers expressed in these areas would be highly informative in this regard. It is possible that GC cells use this PI, or a suite of PIs with similar acyl chain characteristics, for modification and signaling through phosphoinositides. Alternatively, GC B cells, a cellular pool characterized by high proliferation, may produce an abundant and/or diverse pool of PIs as cellular mediators to accommodate relevant proliferation signaling (37). Finally, GC cells may simply increase their PI repertoire to facilitate physical phenotypic changes. Sorted tissue-derived cells demonstrated unique subsets of the similar MSI and extracted whole tonsil lipidomes, further confirming our hypothesis that these cells can be differentiated by their unique lipid signatures serving as molecular ‘barcodes’.

Our data demonstrate a compartmentalization of the human lymphoid organ immune system at different levels including cells, lipids, and secreted protein factors. In line with previously described data (20), this anatomical compartmentalization most likely reflects functional compartmentalization too. We propose an experimental approach of complementary methodologies that can provide unpresented information regarding biological factors contributing to this tissue spatial organization of the immune system. A multimodal tissue analysis integrating the high resolution and volumetric (z axis) analysis of scanning confocal imaging combined with the high throughput and data-rich performance of modern MSI, especially for targets with no known probes, can lead to discovery of novel biomolecules with potential roles for the described immune system anatomy. Furthermore, the current study could serve as a reference for future studies comparing the tissue organization of the human immune system between health and disease.

## Acknowledgements

The presented work was supported by the Intramural Research Program of the Vaccine Research Center, NIAID, NIH, USA.

## Data and materials availability

all data related to this study are present in the main paper and supplementary figures.

## Author Contributions

BLO: Conceptualization, Data curation, Formal Analysis, Investigation, Methodology, Project administration, Visualization, Writing-original draft, Writing-review & editing; JAV-D: Formal Analysis, Investigation, Methodology, Writing-original draft, Writing-review & editing; EM, AM, KM & GF: Formal Analysis, Investigation, Methodology, Visualization, Writing-review & editing; KI: formal analysis, investigation; DA: Investigation, Methodology, Visualization, Writing-review & editing; LDL: Methodology, Resources; ASW: Methodology, Resources, Writing-original draft, Writing-review & editing; RAK: Funding acquisition, Project administration, Resources, Supervision, Writing-review & editing; CP: Conceptualization, Data curation, Formal Analysis, Funding acquisition, Methodology, Project administration, Resources, Supervision, Visualization, Writing-original draft, Writing-review & editing

---

**Supp. Fig. 1. Quantitative analysis of confocal generated images using Histocytometry.** The original image as well as the gating strategy for the identification/quantitation of relevant cell subsets is shown. The histocytometry identified Regions of Interest (ROIs) are also shown.

**Supp. Fig. 2. (A)** General workflow for lipid mass spectrometry imaging with space-directed tandem mass spectrometry structure elucidation scheme. Tissues are thaw mounted onto ITO slides, single stage MSI is performed, ion images are extracted to fill the entire tissue space in a multiple ion image (MII), individual extracted ion images (EII) are used to guide the laser to the coordinates of the ion of interest, and product ion spectra are generated for structure elucidation. **(B)** Generated MS images for three ions in two different tonsil specimens at two different lateral resolutions (50 μm and 20 μm). The same three ions (*m/z* 703, 725, and 728) are plotted in each image on the same scale. **(C)** Example of MS images from the same data set processed in three different ways: No normalization raw data (top), all data normalized to the total ion current (TIC, middle), and all data normalized to the root mean square (RMS, bottom).

**Supp. Fig. 3.** Representative positive ion mode images for ions specific to the follicular area in a human tonsil. 20 ions were detected and given putative identifications based on precursor mass alone. Further MS/MS experiments were used to confirm lipid class for the ions of interest.

**Supp. Fig. 4. (A)** One human tonsil section imaged with negative ion mode MALDI-TOF mass spectrometry at two different lateral resolutions (50 μm and 20 μm). Seven ions of interest were extracted to fill the entire tonsillar tissue area and to represent the known tonsillar anatomy. **(B)** Representative negative ion mode MS images for ions specific to the follicular area in a human tonsil. 24 ions were detected and given putative identifications based on precursor mass alone. Further MS/MS experiments were used to confirm lipid class for the ions of interest.

**Supp. Fig. 5. (A)** Comparison of images of two tonsil sections taken from similar tissue strata using a nuclear stain (JoPro) confocal microscopy (top) and MSI with two ions shown that distinguish the follicle from the extrafollicular area (bottom). (B) Second tonsil replicate MSI with box and whiskers plot demonstrating significantly different (p < 0.002) median expression of three lipids (*m/z* 715, 743, and 797) in the tonsillar follicle versus the extrafollicular area. **(C)** Experimental pipeline for concurrent tonsil cell MALDI-TOF lipidomics using flow cytometry and cell sorting and *in situ* MSI for cryosectioned tonsil tissue.

