## Supplementary figures and images for "Multilevel human secondary lymphoid immune system compartmentalization revealed by complementary multiplexing and mass spectrometry imaging approaches"

### Supplemental Figure 1

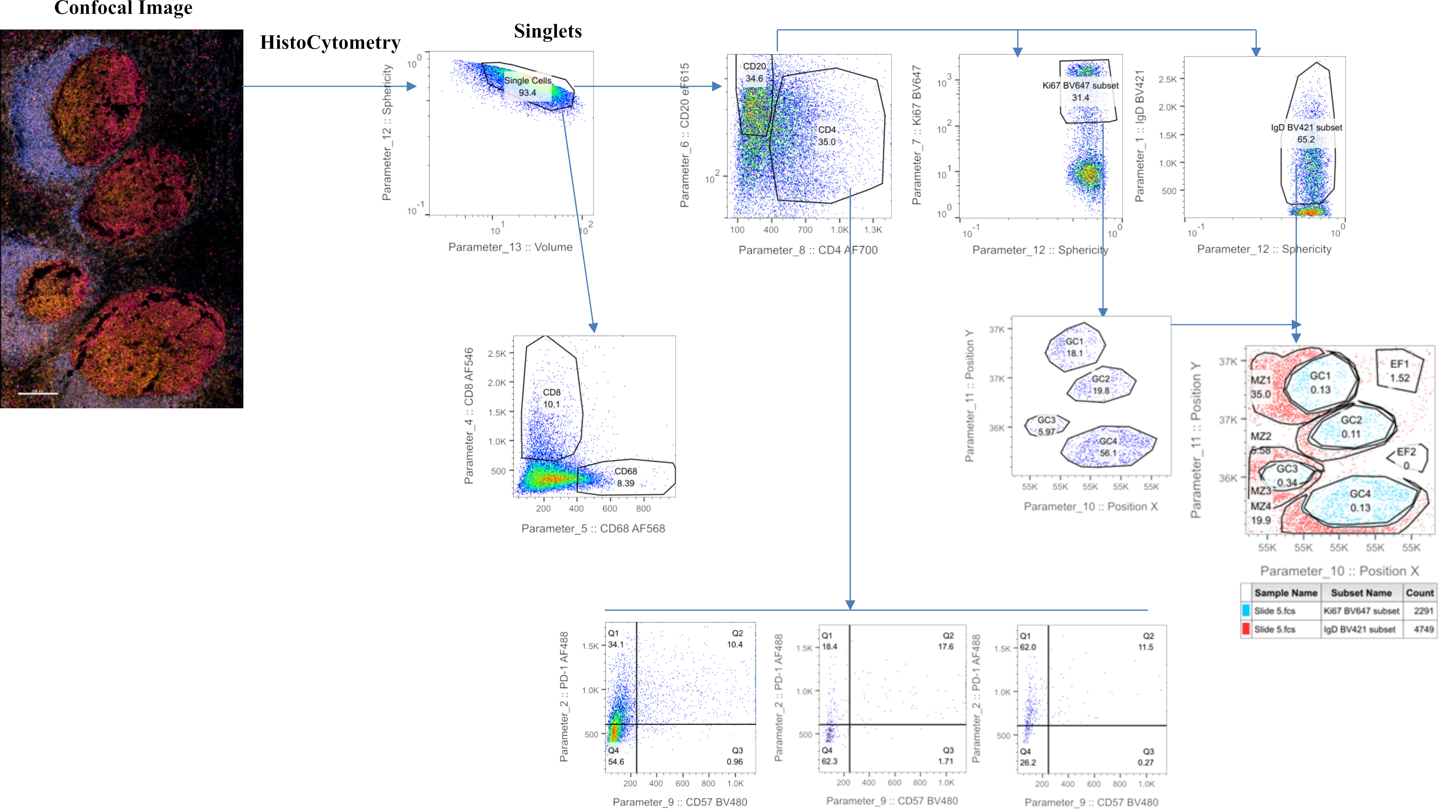

### Supplemental Figure 2

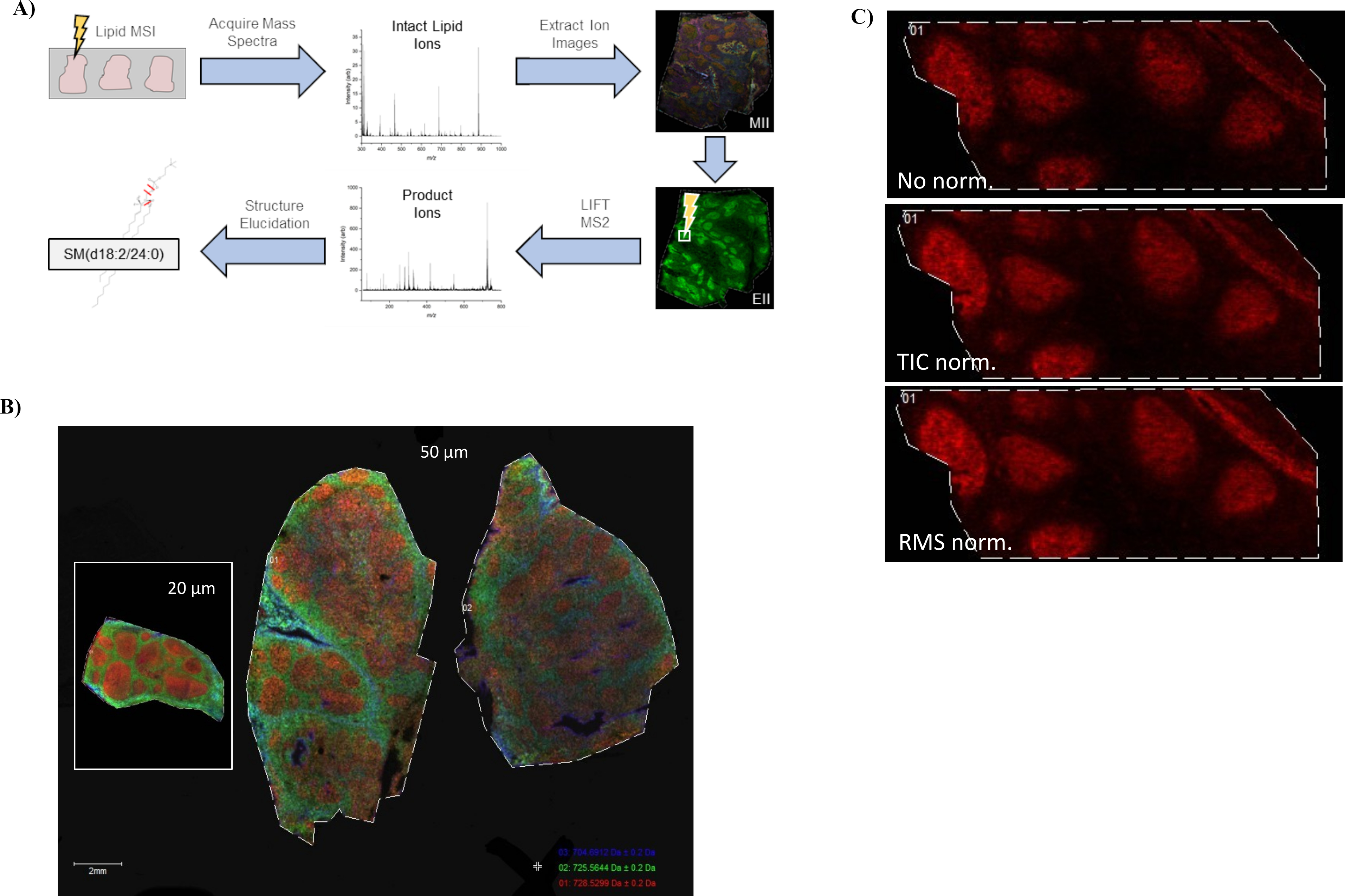

### Supplemental Figure 3

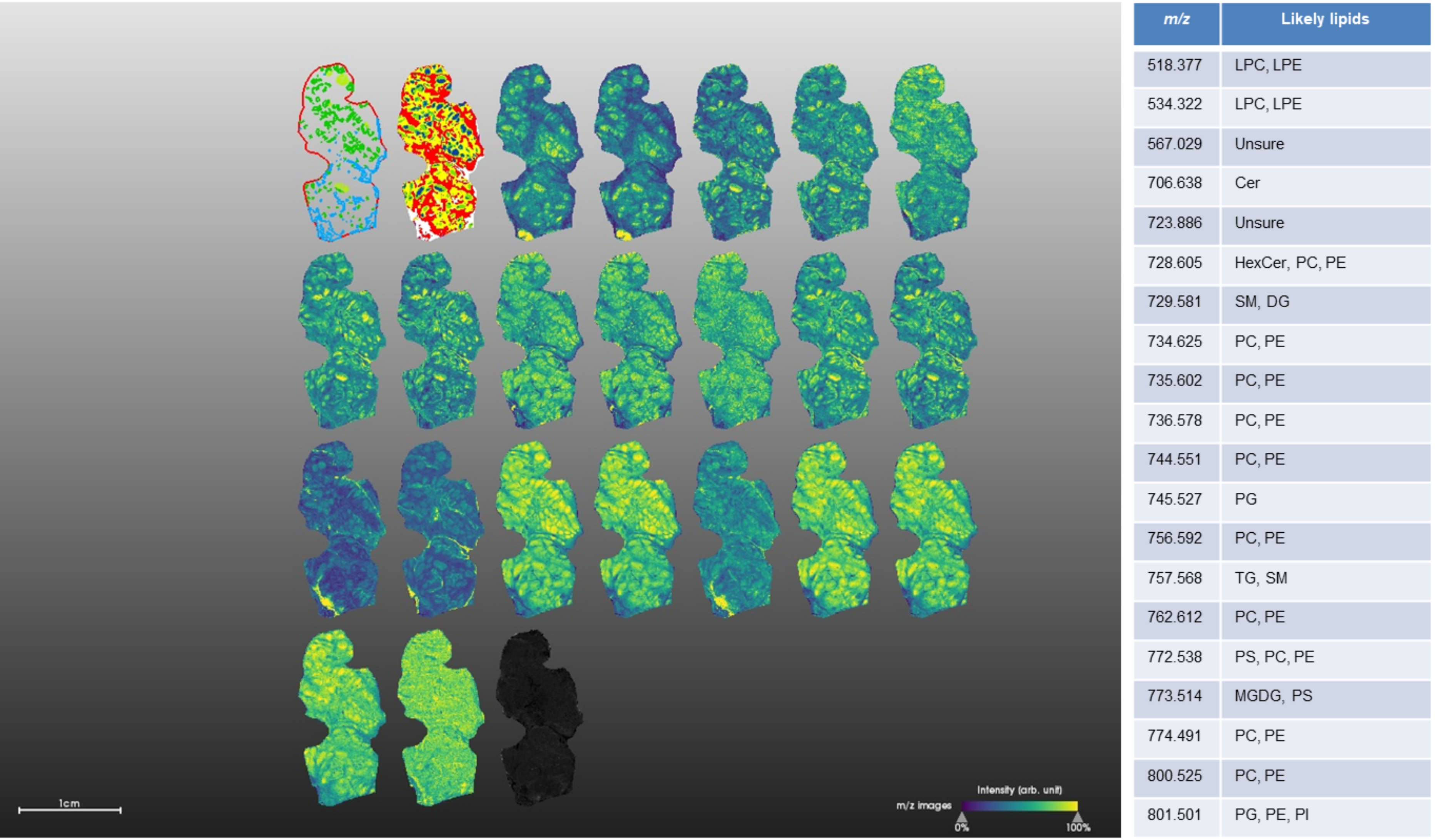

### Supplemental Figure 4

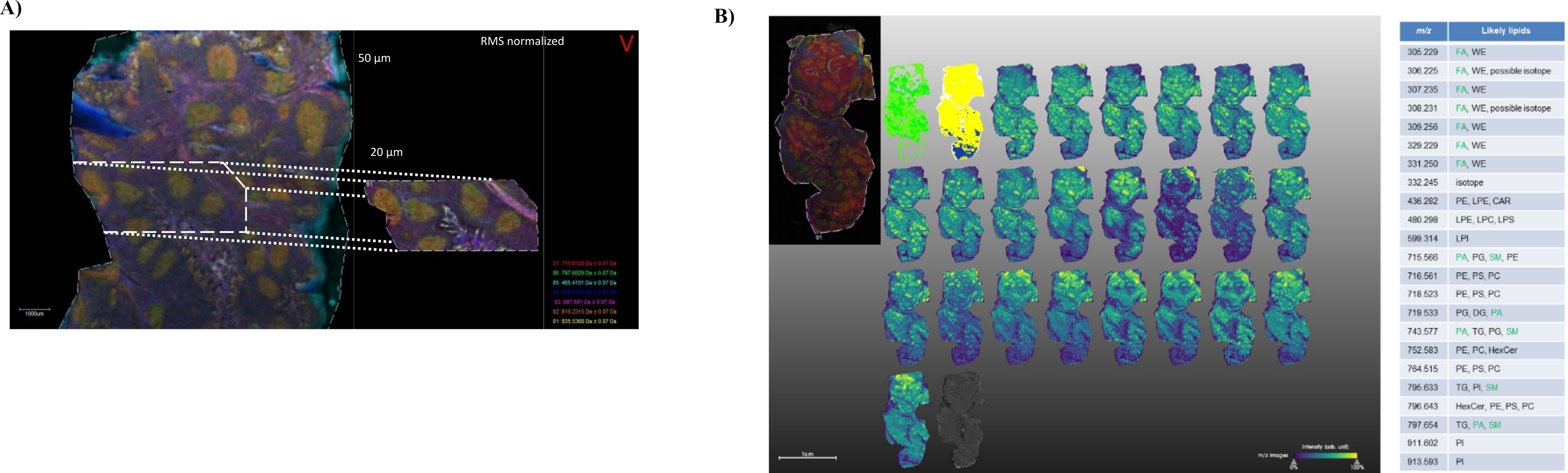

### Supplemental Figure 5

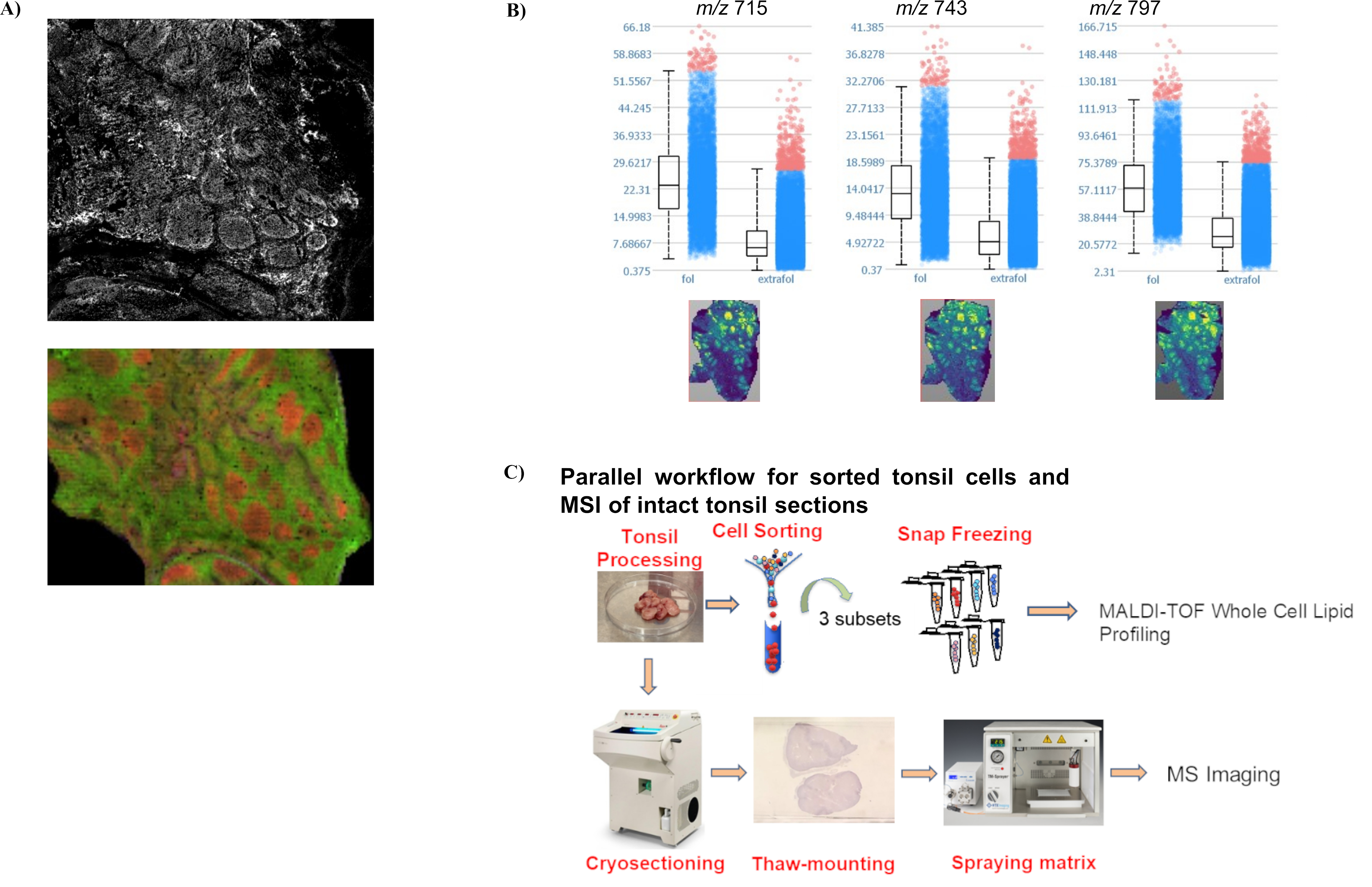
